# Shape analysis of biomimetic and plasma membrane vesicles

**DOI:** 10.1101/2024.10.05.616773

**Authors:** Rajni Kudawla, Harshmeet Kaur, Tanmay Pandey, Tripta Bhatia

**Affiliations:** Department of Physical Sciences, Indian Institute of Science Education and Research, Mohali, Knowledge City, SAS Nagar, Manauli PO 140306

**Keywords:** 3D Visualization, Vesicle morphology, Vesicle shape analysis, Confocal imaging, Point cloud

## Abstract

Giant membrane vesicles (GUVs) and Giant plasma membrane vesicles (GPMVs) are models for examining membrane properties. We attempted to conduct a comparative study of reduced volume of vesicles with different lipid compositions, solution symmetry, solution asymmetry, and membrane charge. The vesicular morphology is studied using three-dimensional visualization techniques. The vesicles may not be exactly similar to spheres, but they may have some fluctuations resulting in changes in their shape. To understand these shape deformations, we visualize them as confocal image stacks. Our experimental observations indicate that the charge of the membrane affects the deflation of the GUVs in the presence of trans-bilayer sugar asymmetries. The lipid bilayers of our GUVs contain a uniform distribution of the lipids in the two leaflets, which implies that the two leaflets have the same lipid composition and there is no asymmetries in this composition. However, they obtain trans-bilayer asymmetries through asymmetric adsorption or desorption layers of different solution compositions adjacent to each leaflet. The deformation of the GPMVs shapes extracted from cells with trans-bilayer buffer asymmetries and the composition asymmetries is also estimated and compared with biomimetic membranes.

## Introduction

Giant unilamellar vesicles (GUVs) and giant plasma membrane vesicles (GPMVs) are two model systems widely used to study membrane properties. GUVs typically have diameters ranging from 10 − 100 *µm* ^[1−4]^. GPMVs also fall within a similar size range, typically between 5 − 20 *µm* in diameter ^[5−7]^. GUVs are spherical vesicles that consist of a single lipid bilayer and are formed using the electroformation^[8]^, gel-assisted method ^[9]^ or, extrusion methods. GPMVs are derived directly from the plasma membrane of cells using the chemical induction method ^[5]^.

The study of vesicle shapes in two-dimensional imaging, which assumes the shape of a vesicle based upon the single-slice cross-sectional contour, opens up the possibility of a vague assumption of a vesicle being axially-symmetric along all axes. The vesicle is an ellipsoid, and the non-deflated vesicles may resemble a sphere, a special case of the ellipsoid, where all three semi-axis are the same. For a single-cut image of a stack, its shape has various possibilities; it can be nearly spherical or have an elongation or compression on one side of the axis perpendicular to the plane of the image, resulting in shape deformation. We need to count on all of the possibilities of vesicle shape; one is confirmed when the vesicle is visualized using the point cloud of the stack of images. When images are captured, the vesicles may show significant motions; we use the affine transformation^[10]^ to align the contours. We employ a three-dimensional visualization technique to understand the vesicle shape and its properties. We tried to conduct a comparative study of the vesicle shape of GUV prepared with different lipid compositions or sugar solution inside and outside of GUV and with GPMV extracted from HEK293 cells.

## Materials and Methods

### Materials

We purchased all lipids from Avanti Polar Lipids in the form of powder. For storage, we dissolve the lipid powder in chloroform and keep it at −20°*C*. We have used 1,2-Dioleoyl-sn-glycero-3-phosphocholine lipids (DOPC) (catalog no. 850375P) and 1,2-Dioleoyl-sn-glycero-3-phospho-(1’-rac-glycerol)(sodium salt) (DOPG) lipids (catalog no. 840475) to make vesicles. DOPC is zwitterionic in nature, whereas DOPG is a negatively charged phospholipid. For making a buffer solution, Sucrose (purchased from Emplura) and glucose purchased from Sigma-Aldrich (CAS no. 50-99-7) were used. For making a buffer solution, milli-Q water is used as a solvent. For preparing GPMVs, we used NaCl (from Sigma-Aldrich, CAS no. 7647-14-5), CaCl_2_ (from Avra, Catalog No. ASC2461), HEPES (from Sigma-Aldrich, H3375), KOH (from SRL, CAS no. 1310-58-3), dithiothreitol (DTT) (CAS no. 3483-12-3, trypsin and formaldehyde (HCHO)(CAS no. 50-00-0) are purchased from Himedia, HEK293 cell line (epithelial cells from the kidney of a human embryo that have adherent properties) purchased from ATCC (CAT no. CRL-1573)

### Preparation of GUVs

The electroformation method is used for the production of giant unilamaller vesicles. In this method, lipids are coated on two glass plates coated with indium tin oxide (ITO) and then choloroform is evaporated in vacuum. Putting the Teflon spacer between two ITO slides coated with lipids, the lipids are facing each other and a chamber is created. The buffer solution was filled into the chamber with a syringe and connected via copper tape to the function generator. Two clamps were used to keep the ITO glass plates together to prevent any solution from escaping the cabinet. An AC sinusoidal field was applied to the electroswelling process. In the electroswelling process, peak voltages (200 − 2000) mV and frequencies (4 − 10) Hz were used in steps.

### Preparation of GPMVs

GPMVs were isolated from the HEK-293 cell line. There are a large number of methods for the extraction of GPMV from adherent cells. In this experiment, the chemical induction method was used ^[5]^. Vesiculation buffer, consisting of 150 *m*M NaCl, 2 *m*M CaCl_2_ and 20 *m*M HEPES, was prepared and the pH was maintained at 7.4 using a 1 M KOH solution. Then, the active vesiculation buffer, consisting of 1.9 *m*M DTT and 27.6 *m*M formaldehyde, was prepared using vesiculation (prepared above) as a solvent on the same day of extraction of GPMVs. Now, to isolate GPMVs, a 25 cm^2^ flask of adherent cells (HEK293) was taken, which was trypsinized one day earlier. The medium was taken out from above the cells and the cells adhered to the surface were washed with 2 *ml* of vesiculation buffer. Finally, 1 *ml* of active vesiculation buffer was added to the flask and kept in the incubator at 37^−^ *C*, gently shaking the flask every 10 minutes for 1 hour 30 minutes.

Osmolality of cytoplasm of cells is about 280-300 mOsm/kg. To calculate the total osmolality of the complete culture media consisting of 89% Gibco RPMI 1640 culture media, 10% FBS, and 1% antibiotic solution, we need to consider the contributions of each component. Thus, the total osmolality of the culture media with 10% FBS, 1% antibiotic solution, and 89% RPMI 1640 is approximately 283.65 mOsm/kg. For GPMVs extraction, cell culture media was removed and we added the active vesiculation buffer containing Vesiculation buffer, consisting of 150 *m*M NaCl, 2 *m*M CaCl_2_ and 20 *m*M HEPES, 1.9 *m*M DTT and 27.6 *m*M formaldehyde (HCHO) giving a total osmolality as 355.5 mOsmol/kg. Therefore, osmolality inside GPMVs is around 280-300 mOsm/kg and osmolality outside GPMVs is around 355.5 mOsmol/kg.

### Phase-contrast microscopy

To observe GUV under a microscope, 20 *µl* GUV were mixed with 30 *µl* of glucose buffer of different concentrations on a coverlip (24 × 60 *mm*). Wait for ten minutes to settle down the GUVs and then start taking observations. For the observation of GUVs and GPMVs, we used phase contrast microscopy (Zeiss Axio Observer 5) equipped with a 20x objective (PL APO 20x/0.8 Ph2 M27). The objective has a working distance of 0.55 *mm* and a field of view of 25 *mm*. A Zeiss Axiocam 506 mono camera was used to record all photos and videos. It gives a two dimensional view of the three-dimensional vesicles. In phase contrast microscopy, observation can only be performed for the equatorial plane of the sample. For three-dimensional shape analysis, confocal microscopy is used.

### Confocal microscopy

For three-dimensional imaging, a confocal microscope (Zeiss Axio observer LSM 980 Airyscan-2) equipped with a PlanApochromat 63x/1.40 oil immersion objective was used. The *z*-stack of GUVs with varying concentration imbalance and GPMVs was taken for the shape analysis. To take the *z*-stack, we must select a *z*-range for our vesicles. In other words, you can find the top and bottom of your sample by focusing across the entire thickness. The size of the *z* step (interval between two slices) is 0.5 *µm*, and the total number of *z* slices depends on the size of the vesicle. The pinhole size used for optimal sectioning is 5 AU (293 *µm*). The scan area (scan zoom) is taken as 2x. For GUVs and GPMVs, Nile red dye was used for the fluorescence. The excitation and emission wavelength for nile red are 559 *nm* and 636 *nm*, respectively.

## Results and Discussion

### Imaging

Three-dimensional images of membrane compartments are obtained by a confocal microscope in the form of stacks, each part of which corresponds to a constant value of *z* and a plane parallel to the *xy* plane. The stacks obtained in this article, are captured as single-channel images ^[11]^.

Each section of the stack captures a cross section of GUVs and GPMVs. We apply a threshold to the image and convert the slices if the grey-scale into binary images. Thresholding helps us to select the contours of the GUV cross-section effectively. Once the contours are identified, we mask them in the original grey scale image. This results in images that contain non-zero pixels only within or on the contours of GUVs and GPMVs.

### Alignment of confocal stacks

Vesicles may undergo significant translation movements, and images are captured. The translation movement of the vesicles is considered a movement between the centre of the stack^[11]^. Since contours are the boundaries of non-zero pixels in the mask image, we use contours to refer to the boundaries of non-zero pixels in the mask image. After masked images are obtained, the overlapping of the vesicles along each other produces non-alignment curves due to the movements of the vesicles. To correct the alignment of the curves, we calculate the translation required for each curve. The geometric centres of the slices are calculated using the masked image moments^[12–14]^.

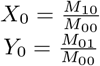

where *M*_*ij*_ are the spatial moments of the image ^[12–14]^. The (*X*_0_, *Y*_0_) is the coordinate of the center of the masked image.

### Lateral shift calculation

The lower part of the stack is used as a reference curve and the center of reference coordinate (*x*_*ref*_, *y*_*ref*_) is obtained from the moment of the lower masked part. In order to calculate the translation matrix for each slice, we calculate the (Δ*x*, Δ*y*) shift in (*x*_*i*_, *y*_*i*_) as a transfer between the center of the slice and the center of the reference slice.

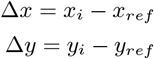

### Affine transformation

The image is aligned by affine transformation^[15]^. Affine conversion is a linear mapping method that rotates, converts, and scales images. The affine transformation matrix *A* of a two-dimensional image can be represented as follows

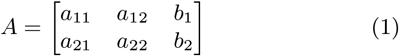

The translation matrix *T* and the rotation matrix *R* can be written as

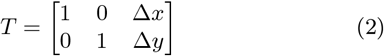

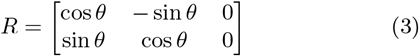

So, the affine matrix A can be represented as

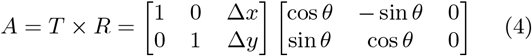

So, we can write the affine transformation matrix as,

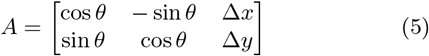

Since we account for only the translational shift, not the rotational shift, we put *ω* = 0. Therefore, we get

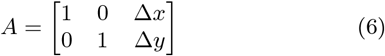

The affine transformation matrix *A* is applied to the slices of the *z*-stacks, and the aligned images are obtained.

### Three dimensional projection

A two-dimensional image is a set of pixels. The pixel co-ordinates are represented as pairs (*x, y*), *x* represents the column index, *y* represents the row index ^[16]^. The source of the coordinate system is (0, 0), in the upper left corner of the image, and the positive axis *x* is extended horizontally to the right, and the positive axis *y* is extended vertically downwards ^[17]^. Since each slice of the *z* stack corresponds to a certain *z* value, the pixel of the given 2-dimensional (*x, y*) slice has the same *z* coordinate. Thus, each pixel has a co-ordinate (*x, y, z*), which can be displayed in a 3D coordinate space.

### Shape deformation, Reduced Volume (*v*)

After a three-dimensional projection, the surface area, volume and reduced volume of the vesicle can be calculated. The surface of the vesicle is *S* and the volume of the vesicle is *V*. The volume of the vesicle *V* attains its maximal value when the vesicle has a spherical shape. The reduced volume (*v*)^[**?**]^ of the vesicle is defined as

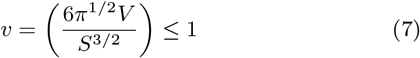

Table 1 shows the average value of the reduced volume (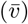) with the error bar (Δ*v*) of the GUV samples in equilibrium with a solution of 300 *m*M sugar concentration, inside and outside. The differences in concentration throughout the GUV membrane are absent. The data-based graph is shown in Fig. 3. Solution asymmetries are present in samples I, II, i.e. sucrose is present inside the GUV and glucose in the exterior compartment of GUVs. In addition, the II-IV samples consists of charged lipid membranes.

**Table 1.** The average value of the reduced volume (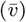) with the error bar (Δ*v*) of flUVs samples composed of different lipid(s) in equilibrium with 300 *m*M solution inside-out. The plot with data is shown in Fig. 3. The solution asymmetry is present for samples I, II. Samples II-IV are composed of charged lipid membranes.

| Sample no. | membrane, in/out (300 mM) solution | $\bar{v} \pm \Delta v$ |
| --- | --- | --- |
| I | DOPC, suc/glu | $0.89 \pm 0.08$ |
| II | DOPC: DOPG (10%), suc/glu | $0.56 \pm 0.02$ |
| III | DOPC: DOPG (10%), suc/suc | $0.79 \pm 0.08$ |
| IV | GPMVs, buffer/buffer | $0.85 \pm 0.08$ |
| V | DOPC, suc/suc | $0.98 \pm 0.02$ |

**Figure 1.**
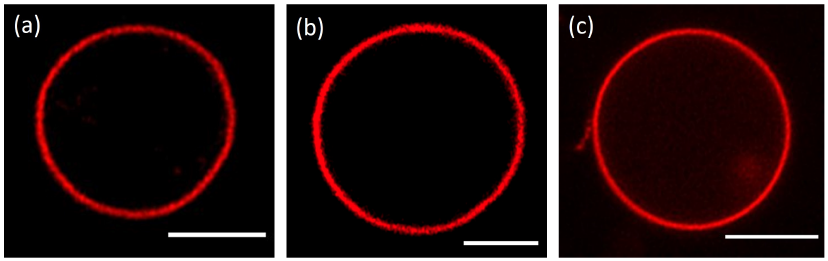
Confocal fluorescence image of vesicles. **(a)** DOPC flUV in 300*mM* Sucrose, **(b)** DOPC:DOPfl (10%) flUV in 300*mM* Sucrose, **(c)** flPMV extracted from HEK293 cell line. The scale bar is 5 *µm* applied to all the images.

**Figure 2.**
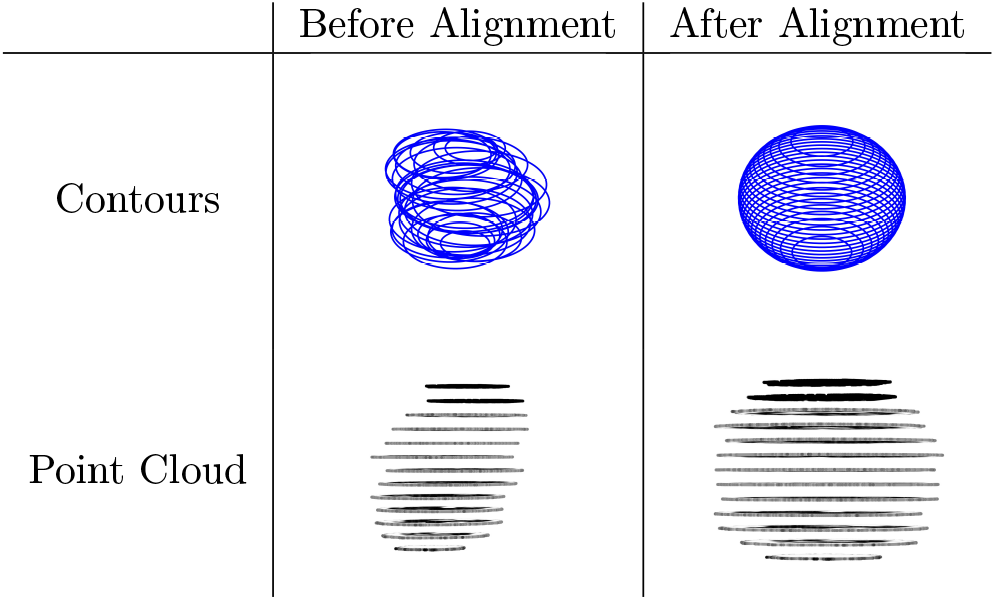
The contours of the slices are translated using the shift calculated by applying an affine transformation matrix so the centre of the contours are aligned on a common axis for the complete stack.

**Figure 3.**
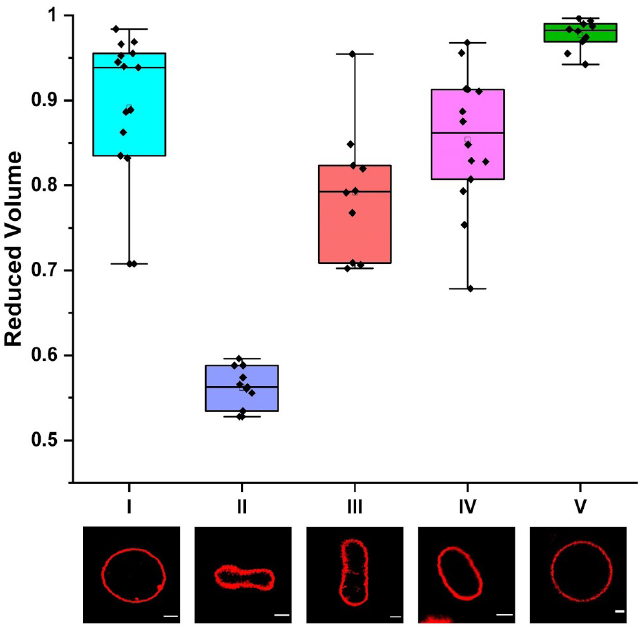
Comparison plot for the reduced volume of the GUVs samples. **I**: DOPC GUVs (300 *m*M sucrose inside and 300 *m*M glucose outside), **II**: DOPC:DOPG (10%) (300 *m*M sucrose inside and 300 *m*M glucose outside) GUVs, **III**: DOPC:DOPG(10%) (300 *m*M sucrose inside and outside) GUVs, **IV**: GPMVs, **V**: DOPC GUVs (300 *m*M sucrose inside and outside). On the *X* axis, the sample number is displayed. The images show the *XY* snapshot of the GUV equilibrium shape for the sample number indicated. The red colour is the GUV membrane. The scale bar applies 5 *µm* to all images. The data is shown in Table 1.

### Shape deflation of membrane vesicles

To understand the deflation, we prepare GUVs with two different lipid compositions, DOPC and a mixture of two lipids DOPC:DOPG (10%) in 300 *m*M sucrose solutions.

#### Role of solution asymmetry in shape deflation

GUVs were prepared in 300 *m*M sucrose inside the GUV and were transferred in the same concentrations of glucose outside the GUV thereby generating a sugar solution asymmetry across the membrane. In our system, the membrane shape change is generated by the asymmetric adsorption/desorption layers formed by the sugar molecules adjacent to the two leaflets. Because these two layers involve different types of sugars, the bilayers become asymmetric. ^[1–3,18–21]^

To understand deflation of shape, we prepare GUVs with two different lipid compositions, DOPC and a mixture of two lipids DOPC:DOPG (10%). The lipids of DOPC GUV membrane and DOPC/DOPG GUV membrane contain only a uniform lipid composition in the bilayer, implying that the two leaflets have the same lipid composition and there is no asymmetries from this point of view. However, they acquire transbilayer asymmetry through the asymmetric adsorption/desorption of sugars adjacent to the two leaflets. Compared to sucrose, glucose have additional chemical groups that can come into contact with the lipids in the external leaflet and glucose is smaller than sucrose. Therefore, glucose can form more densely packed adsorption layers at the outer leaflet. ^[1–3,18]^ The plot with the data is shown in Fig. 3 and given in the Table 1.

Fig. 3 shows shape deflation in the absence of sugar concentration imbalance for; DOPC GUVs (sample V) with solution symmetry, GPMVs extracted from the cells (sample IV) with solution asymmetry, DOPC: DOPG (10%) (sample III) membrane with solution symmetry, DOPC: DOPG (10%) (sample II) membrane with solution asymmetry, and for DOPC GUVs (sample I) with solution asymmetry. osmolality inside GPMVs is around 280-300 mOsm/kg and osmolality outside GPMVs is around 355.5 mOsmol/kg.

GUVs enable the passage of small uncharged molecules such as H_2_0, O_2_ and CO_2_, as well as H_3_O^+^ and OH^−^, and prohibit the passage of other ions or larger water-soluble molecules such as glucose^[22]^. Thus, these solvents act as osmotic active particles and generate osmotic pressures on vesicular membranes. The concentrations of solvents within and outside the vesicle determine these pressures. For Table 1 and in Fig. 3, the concentration imbalance is absent giving the osmotic pressures on vesicular membranes as the same from inside-out. The reduction in DOPC GUV volume in the absence of concentration imbalance across the membrane is 0.89 *±* 0.08, or 3% from a spherical shape, however for DOPC: DOPG (10%) GUVs it is 0.56 *±* 0.02, or around 50% from the spherical shape, suggesting a role of the membrane charge, discussed below.

#### Role of membrane charge in shape deflation

The values of reduced volume for charged GPMVs and for DOPC/DOPG vesicles are much less compared to 50% reduction in the spherical volume of DOPC: DOPG (10%) GUVs in the presence of sugar asymmetry with no concentration imbalance, shown in Table 1.

DOPC is a zwitterionic lipid, while DOPG is a negatively charged lipid and GPMVs derived from cells are charged plasma membranes. GUVs membrane composed of DOPC and DOPG (10%) has a negative overall charge on the lipid layer, while DOPC GUVs do not have an overall charge on the lipid layer. GPMVs derived from the plasma membrane of the cell have the same buffer medium inside and outside, across the membrane. In the absence of concentration imbalance of sugars, buffers across the membrane, the GUVs membrane composed of DOPC, have on the average value of reduced volume as 0.89 *±* 0.08 which is a larger value than 0.56 *±* 0.02 for vesicles composed of charged lipid bilayer, DOPC: DOPG (10%) respectively in the presence of sugar asymmetry, i.e. sucrose inside the GUV and glucose in the exterior compartment of GUVs.

In order to understand the role of membrane charge only, we have performed experiments in the absence of the sugar asymmetry, i.e. sucrose is present inside the GUV and in the exterior compartment of GUVs. The reduction in DOPC: DOPG (10%) GUV volume (sample III) in the absence of sugar asymmetry and concentration imbalance across the membrane is 0.79 *±* 0.08, or a maximum of 13% from a spherical shape. The reduction in GPMVs volume (sample IV) in the presence of solution asymmetry and no concentration imbalance across the membrane is 0.85 *±* 0.08, or a maximum of 7% from a spherical shape, in equilibrium.

We find that the DOPC: DOPG (10%) GUVs membrane show a relatively less reduction in their volume compared to the case when the solution asymmetry is present, shown in Fig. 3 and in Table 2. Our experimental observation suggests that the introduction of the overall charge into the lipid bilayer assists in deflation of GUVs shape for the same concentration imbalance of the solutes across the membrane, in the presence of sugar asymmetry. We find the shape deflation for GPMV derived from cells as 0.85 *±* 0.08, or a maximum of 7% from a spherical shape.

**Table 2.** The comparison of the percentage change of the reduced volume (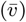) of vesicles in equilibrium in the absence of the concentration imbalance (Δ*C* = 0) across the GUV membrane. The solution asymmetry is present for samples I, II, and is absent for samples III-V.

| Negatively charged membrane, DOPC/DOPG |  |  |  |
| --- | --- | --- | --- |
| Sample no. | Inside | Outside | $\Delta v$ |
| II | Sucrose | Glucose | 50% |
| III | Sucrose | Sucrose | 13% |
| Neutral membrane, DOPC |  |  |  |
| Sample no. | Inside | Outside | $\Delta v$ |
| I | Sucrose | Glucose | 3% |
| V | Sucrose | Sucrose | < 1 % |
| GPMVs derived from cell |  |  |  |
| Sample no. | Inside | Outside | $\Delta v$ |
| IV | Buffer | Buffer | 7% |

#### Role of concentration imbalance in shape deflation

We prepare GUVs with two different lipid compositions, DOPCs and DOPC with DOPG (10%) in 300*m*M of sucrose. GUVs were transferred in relatively higher glucose concentrations outside the GUV to achieve a concentration difference across the membrane at *t* = 0, shown in Table 3. The GUVs were transferred in solution with osmolarity differences of 50 *m*M, 100 *m*M and 150 *m*M,, shown in Table 3 and the respective reduced volume is calculated, shown in Table 4. The solution asymmetry is present for samples I, II. Fig. 4 and Fig. 5 show reduced volume plot for samples I, II with an increasing concentration imbalance across the membrane.

**Table 3.** The equilibrium value of the solution concentration (*C*) of GUVs samples. The first column shows the sucrose concentration (*C*_i_) used for GUV preparation. The second column shows the glucose concentration (*C*_o_) in which GUV are transferred for imaging. Differences in inside-out concentration of sugars is (Δ*C*) shown in the third column. *C*_eq_ is the equilibrium concentration of sugar solution inside-out, after osmotic equilibrium across the GUV membrane is attained.

| GUVs at $t = 0$ | | | GUVs in osmotic eq. |
| --- | --- | --- | --- |
| $C_i$ | $C_o$ | $\Delta C$ | $C_{eq}$ |
| (mM) | (mM) | (mM) | (mM) |
| 300 | 300 | 0 | 300 |
| 300 | 350 | 50 | 330 |
| 300 | 400 | 100 | 360 |
| 300 | 450 | 150 | 390 |
| GPMVs at $t = 0$ | | | GPMVs in osmotic eq. |
| 280-300 | 355.5 | $\geq 55.5$ | 323 |

**Table 4.** The average value of the reduced volume (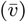) of GUVs samples composed of DOPC (sample I) and DOPC: DOPG (10%) (sample II) lipid(s) in equilibrium. The first column shows the concentration differences (Δ*C*) across the GUV membrane at the time of transfer of GUVs prepared in sucrose and transferred in glucose. The plot with data is shown in Fig. 4 for DOPC membrane and in Fig. 5 for DOPC:DOPG (10%) membrane respectively. The solution asymmetry is present for both the samples I, II.

| $\Delta C$ | $\bar{v}$ | |
| --- | --- | --- |
| (mM) | sample I | sample II |
| 0 | $0.89 \pm 0.08$ | $0.56 \pm 0.02$ |
| 50 | $0.77 \pm 0.09$ | $0.53 \pm 0.05$ |
| 100 | $0.74 \pm 0.10$ | $0.51 \pm 0.05$ |
| 150 | $0.71 \pm 0.05$ | $0.51 \pm 0.02$ |

**Table 5.** Eccentricity values of DOPC:DOPG (10%) [Sucrose and Glucose] GUVs with no concentration imbalance. The X-Y plane is the image plane whose cross-sectional image has been captured. The z-axis is perpendicular to the image plane, which we obtain through three-dimensional modelling of the vesicle.

| GUV no. | Eccentricity YZ | Eccentricity XY |
| --- | --- | --- |
| 1 | 0.7871 | 0.1729 |
| 2 | 0.7821 | 0.0840 |
| 3 | 0.7836 | 0.2696 |
| 4 | 0.7940 | 0.0310 |
| 5 | 0.7844 | 0.1286 |
| 6 | 0.7871 | 0.0420 |
| 7 | 0.7840 | 0.0349 |
| 8 | 0.7846 | 0.1305 |
| 9 | 0.7800 | 0.0241 |
| 10 | 0.7877 | 0.2274 |
| 11 | 0.778 | 0.05706 |

**Figure 4.**
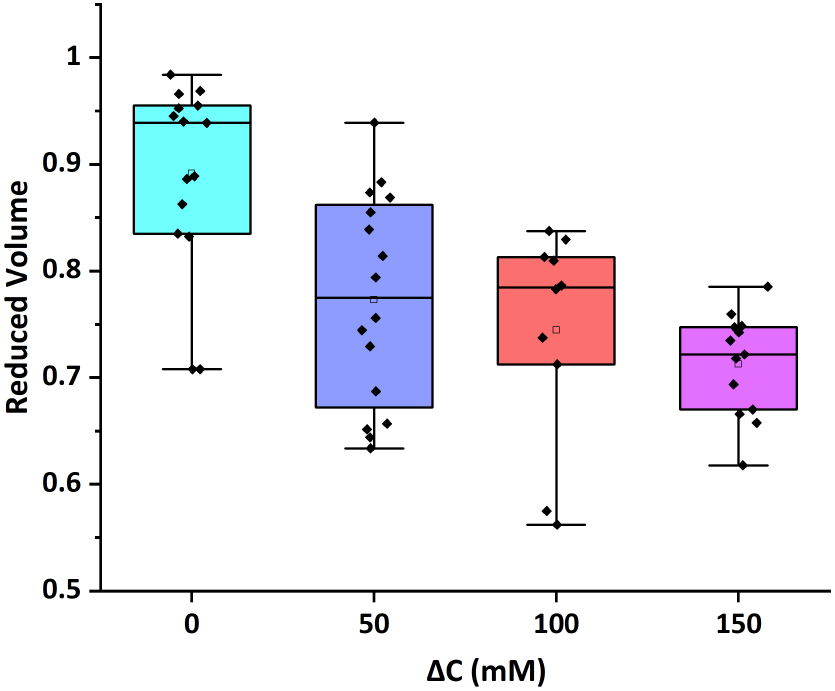
The box plot shows a reduction in the volume of DOPC lipid GUVs in the presence of sugar concentration differences. The *x*-axis shows the difference in the concentration of Sucrose and Glucose solutions. All GUVs are prepared in 300 *mM* Sucrose solution. The concentration of Glucose solutions is 300 *m*M, 350 *m*M, 400 *m*M, and 450 *m*M, respectively, which gives the difference in the concentration as (0, 50, 100, 150) *m*M shown along the *x*-axis.

**Figure 5.**
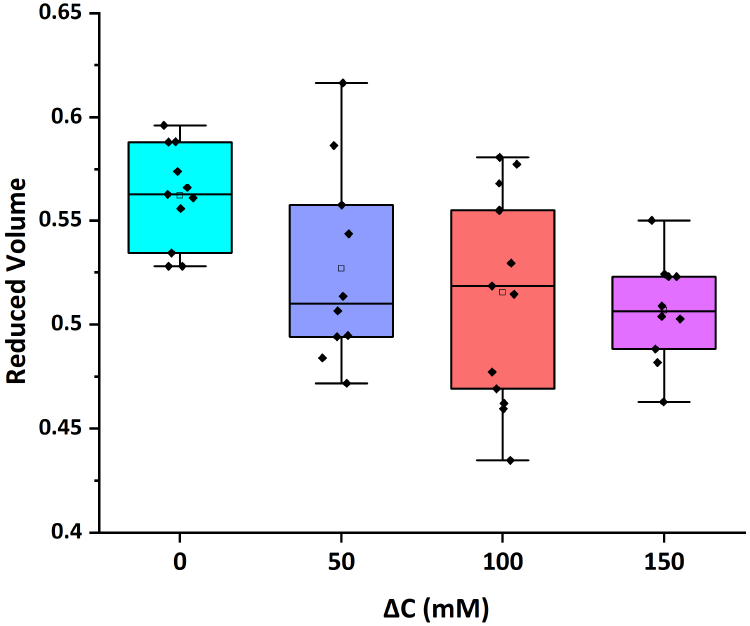
The box plot shows a reduction in the volume of DOPC:DOPG (10%) lipid GUVs in the presence of sugar concentration imbalance. The *x*-axis shows the difference in the concentration of Sucrose and Glucose solutions. All GUVs are prepared in 300 *mM* Sucrose solution. The concentration of Glucose solutions is 300 *mM*, 350 *mM*, 400 *mM*, and 450 *mM* respectively, which gives the difference in the concentration as (0, 50, 100, 150) *mM* shown along the *x*-axis.

As observed in Fig. 4 and Fig. 5, with increasing concentration imbalance across the GUV membrane, the GUV volume reduces, so the vesicle deflates more with the increasing concentration imbalance across the membrane. In response to concentration imbalance, i.e, different values of *C*_i_ and *C*_o_ in Table 3, the GUV expels water from the interior to exterior solution, thereby reducing its volume by osmotic deflation. However, the quantitative analysis described in the main part of our manuscript applies to the equilibrium shapes that the GUVs attain after reaching the osmotic equilibrium. *C*_eq_ in Table 3 is the equilibrium concentration of sugar solution inside-out, after osmotic equilibrium across the GUV membrane is attained.

From Table 4, we can observe that the deflation of DOPC GUVs is relatively higher for the increasing concentration imbalance values in the range (0 − 150) *m*M in the presence of sugar asymmetry, i.e. sucrose inside the GUV and glucose in the exterior compartment of GUVs. The deflation of charged GUVs is relatively less with an increase in the sugar concentration imbalance of (0 − 150) *m*M in the presence of the solution asymmetry across the membrane.

If there is a difference in the concentration of the solvent within relative to outside the vesicle, the dissimilarity of the osmotic pressure moves water through the membrane towards the highest concentration of the solvent ^[22]^. If the concentration of solution outside the vesicle is higher, the water flows out and the vesicle shrinks in volume, resulting in deflation. When the osmolarity difference is higher, if the external solvent concentration is higher, the vesicle decreases in volume until the solvent concentrations are balanced and the osmotic pressure difference is zero at the end. If *C*_eq_ is the equilibrium sugar concentration then Table 3 shows the equilibrium concentration for the samples I and II.

Our experimental observation of the decrease in reduced volume with an increase in concentration imbalance verifies that the extent of deflation is directly proportional to the magnitude of the concentration difference, Δ*C*. The greater the difference, the more water will be driven out, leading to greater deflation for a given membrane composition, shown in Table 3.

The data in Table 2 suggests that those with different sugars inside and outside the GUV are more deflated than those with the same sugar inside and outside, which are nearly spherical. The presence of different sugars inside and outside the GUV can lead to variations in the distribution of water molecules and ions around the lipid bilayer, potentially altering its mechanical properties and response to changes in osmolarity. However, we can note that the reduced volume of DOPC is still higher than both DOPC:DOPG (10%) [Sucrose only] and DOPC:DOPG (10%) [Sucrose Inside and Glucose Outside].

### Shape eccentricity and symmetry

The vesicle shape determination through its two-dimensional cross-sectional image of a vesicle possesses the assumption of being axial-symmetric, which is a very vague assumption to work on. The vesicles are ellipsoids and the spherical vesicles are a special case of ellipsoid vesicles.

Let the side-length of the GUV above its equatorial slice on the axis perpendicular to the plane of the slice be *c*_1_ and below its equatorial slice be *c*_2_. So, for spherical vesicles,

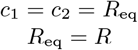

So, for the spherical case, the eccentricity of the vesicle in any plane in the three dimensional space would be zero. If the vesicles have shape variations (deformations or deflation), the eccentricity diverges from zero and becomes different for different planes in the three dimensional space. When we look at the eccentricity of the slice contour from the stack, we get information about the circularity of that cross-section of the GUVs, but we do not get complete details about the vesicle shape. The vesicle may have an oblate or prolate shape or a deflated spherical shape (deformation).

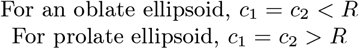

The eccentricity in the *xy* plane was calculated from the contour of the equatorial slice in the confocal stack. From the three-dimensional projection, we obtain the eccentricity of the equatorial cross-section in the *yz* plane.

The eccentricities of the *xy* plane suggest that the cross-section is highly circular, and the vesicle shape, according to the eccentricity of the slice in the *xy* plane, assuming the vesicle to be axial-symmetric, will be highly spherical. After the three-dimensional projection and calculation of reduced volume and eccentricity in a plane perpendicular to the *xy* plane (the *yz* plane), the eccentricity of the cross-section in that plane suggests that the cross-section is very distorted and not even close to being a circle. If both of the cross-sections were nearly circular (i.e., their eccentricity ≈ 0), then the GUV would have been assumed to be nearly spherical. Still, the eccentricity in the *yz* plane suggests the cross-section to be non-circular. So, the vesicle shape is not spherical; it is a deflated sphere (deflated ellipsoid). In such cases,

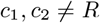

The vesicle may have *c*_1_ = *c*_2_ or *c*_1_ ≠ *c*_2_, the first one resulting in a symmetry along the *xy* plane, and the latter suggesting no symmetry. The ratio 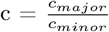 (where, *c*_*major*_ = *max*(*c*_1_, *c*_2_), *c*_*minor*_ = *min*(*c*_1_, *c*_2_)) tells us how much the vesicle is elongated on one side of the axis perpendicular to the image plane. If c = 1, then *c*_1_ = *c*_2_, we could compare the contour radius of the equatorial image plane (R) with *c*_1_ and *c*_2_ to discuss the structure of the vesicle.

## Conclusion

The neutral GUVs samples were prepared with DOPC, one with sucrose solution inside and outside the GUVs (Sample V) and the other with sucrose solution inside and glucose solution outside the GUVs (Sample I). The charged GUVs samples were prepared with DOPC: DOPG (10%) lipid composition, one sample with sucrose solution inside and outside the GUVs (sample II), and the other with sucrose solution inside and glucose solution outside the GUV (sample III). GPMVs are derived from the plasma membrane of the cell have the buffer medium inside and outside, across the membrane (sample IV). The asymmetry between the lipid compositions of the two leaflets give rise to trans-bilayer asymmetry. There are many other ways to generate bilayer asymmetry by many different molecular mechanisms. ^[1,18]^ In this paper, bilayer asymmetry in GUVs is generated by solution asymmetry in which the two leaflets of the bilayers are exposed to two different sugar solutions. Previously, in this case, with sugar solution asymmetry for the bilayers we have observed multispherical shapes as well. ^[2,3]^

In this paper, we validate that the shape of the vesicles depends heavily on the membrane charge, solution asymmetry and solution concentration imbalance. Previously, membranes consisting of charged lipids have been studied. ^[23–25]^ In this study, we observed the changes in the shape of the vesicles of charged GUVs and GPMVs. We found that with the charged and uncharged lipid compositions of GUVs, the vesicles shape deflation depend on the solution asymmetry at constant solute concentration, which emphasize the role of membrane asymmetry and membrane charge in deflation. Furthermore, with the increase in the solution concentration imbalance across the membrane for the same lipid composition, the GUV deflation increases, causing greater shape changes of the GUVs at higher concentration differences, except for DOPC/DOPG (10%) membranes with sugar solution asymmetry. This emphasize the coupling of membrane asymmetry and membrane charge in deflation of GUVs with concentration imbalance and without.

Fig 3 shows the change in the shape of the vesicles and their reduced volume in the absence and presence of membrane asymmetry without concentration imbalance across the membrane. For a given concentration difference of solutions across the membrane to reduce the GUV volume, the shape of the vesicle, for DOPC: DOPG (10%) GUVs is very dissimilar in the presence of sugar asymmetry and without. GUVs with different sugars inside and outside the GUV were more deflated, and their volume reduction is larger compared to GUVs with the same sugar inside and outside the GUV. For DOPC GUV and DOPC: DOPG(10%) (sugar and glucose), Fig 4 and 5 show that when the volume decreases, vesicles deflate more with an increasing solution concentration imbalance. We also studied the GPMVs prepared by HEK293 cells; the resulting reduced volume is found to be lower than DOPC GUVs with and without membrane asymmetry. It is important to note that GPMVs have membrane asymmetry due to compositional heterogeneity and solution trans-bilayer asymmetry both and these are derived from the cellular plasma membranes.

The two-dimensional image (the *XY* cross-section of the vesicle) does not reveal the morphology of the vesicle on the plane perpendicular to the image plane (axis z). To determine the shape of the vesicles, information is also needed about the shape of the vesicles along the z-axis. Suppose we capture the cross section of the vesicle on its equatorial plane to determine its structure. In this case, we must take into account the vague assumption that the vesicle is axially symmetric and base the vesicle morphology on the information contained in the cross-section imaging, which may not be the case exactly. Our model provide a useful way to calculate the reduced volume for membrane compartments and will be shared publicly.

This work allows for a more in-depth understanding of how the complexity of cell membranes composed of lipids and proteins can stabilize curvature and contribute to the classification of shapes by remodelling the membrane charge and membrane asymmetry.

## Author Contribution

RK performed the GUVs experiments, HK performed GPMVs experiments, and TP wrote the code for the reduced volume analysis, performed data analysis. TB designed the experiments and the code. TP, RK, HK and TB wrote the manuscript. All authors discussed the manuscript and commented on it.

## Acknowledgements

This research was conducted within the Indian Institute of Science Education and Research Mohali and supported by the Ramalingaswami grant of DBT (grant DBT/RLF/Re-entry/06/2020).

## Conflict of Interest

All authors have no conflict of interest.

## Notes

### Competing Interest Statement

The authors have declared no competing interest.

## References

[1] R. Lipowsky, Understanding giant vesicles – a theoretical perspective - Reinhard Lipowsky, in The Giant Vesicle Book, CRC Press, Boca Raton 2019.

[2] T. Bhatia, S. Christ, J. Steinkühler, R. Dimova, R. Lipowsky, Soft Matter 2020, 16, 1246.

[3] A. Nowbagh, A. Deshwal, M. Kadu, A. Chaudhuri, S. Maiti, R. Lipowsky, T. Bhatia, ChemSystemsChem 2022, 5, e202200027.

[4] T. Bhatia, J. Membr. Biol. 2022, 255, 637.

[5] Z. Gerstle, R. Desai, S. L. Veatch, Giant plasma membrane vesicles: An experimental tool for probing the effects of drugs and other conditions on membrane domain stability, in Methods in enzymology, volume 603, pages 129–150, Elsevier 2018.

[6] E. Sezgin, H. J. Kaiser, T. Baumgart, P. Schwille, K. Simons, I. Levental, Journal of Visualized Experiments 2012, 63, e3963.

[7] E. Sezgin, H.-J. Kaiser, T. Baumgart, P. Schwille, K. Simons, I. Levental, Nature protocols 2012, 7, 1042.

[8] Z. Boban, I. Mardešić, W. K. Subczynski, M. Raguz, Membranes 2021, 11.

[9] A. Weinberger, F.-C. Tsai, G. H. Koenderink, T. F. Schmidt, R. Itri, W. Meier, T. Schmatko, A. Schröder, C. Marques, Biophysical Journal 2013, 105, 154.

[10] R. Hartley, A. Zisserman, Multiple View Geometry in Computer Vision, Cambridge University Press 2004.

[11] P. Husen, M. Fidorra, S. Härtel, L. A. Bagatolli, J. H. Ipsen, Biophysical Journal 2012, 103, 2304.

[12] M.-K. Hu, IRE transactions on information theory 1962, 8, 179.

[13] J. Flusser, T. Suk, Pattern recognition 1993, 26, 167.

[14] R. Mukundan, K. Ramakrishnan, Moment functions in image analysis: theory and applications, World scientific 1998.

[15] S. Cao, Y. Yu, H. Guan, D. Peng, W. Yan, Remote Sensing 2019, 11, 1708.

[16] R. C. Gonzalez, R. E. Woods, Digital Image Processing, Prentice Hall, 2nd edition 2002.

[17] R. Szeliski, Computer Vision: Algorithms and Applications, Springer 2010.

[18] R. Lipowsky, Advanced Biology 2022, 6, 2101020.

[19] J. E. Rothman, J. Lenard, Science 1977, 195, 743.

[20] H. F. Lodish, J. E. Rothman, Scientific American 1979, 240, 48.

[21] H. Kumar, S. Mandal, R. Yadav, S. Gupta, H. Meena, M. Kadu, R. Kudawla, P. Sharma, I. P. Kaur, S. Maiti, J. H. Ipsen, T. Bhatia, Chemistry and Physics of Lipids 2024, 259, 105374.

[22] H. Wennerström, E. Sparr, J. Stenhammar, Physical Review E 2022, 106, 064607.

[23] A. Fogden, D. J. Mitchell,, B. W. Ninham, Langmuir 1990, 6, 159.

[24] H. N. W. Lekkerkerker, Physica A: Statistical Mechanics and its Applications 1989, 159, 319.

[25] A. C. Rowat, P. L. Hansen, J. H. Ipsen, EPL 2004, 67, 144.

